# Cell-Lens: Advancing LLM Reasoning at Single-Cell Resolution

**DOI:** 10.64898/2026.09.19.752842

**Authors:** Yanchao Li, Ben Gao, Wanhao Liu, Jiaqing Xie, T. Y. Tsui, Yuanyuan Zhang, Yanbo Wang, Tianfan Fu, Yuqiang Li

## Abstract

Recent work applies large language models (LLMs) to single-cell data, but the cell usually reaches the model as a short ranked list of genes. This list drops much of what defines a cell, so the model reasons from a partial view. Yet the information is not lost in the measurement, only in the text. Here, we introduce Cell-Lens, a training-free structured representation of the measured cell. It writes the cell as typed blocks, from paired measurements to marker-supported programs. On CellVerse, SOAR, CellPuzzles, and SC-Arena, across five model families, Cell-Lens improves performance by up to 27.1 points. It also narrows the model gap and reduces API-reported reasoning-token usage. The largest gains occur when a defining RNA marker falls out of the ranked list. In these cases, the paired protein data still capture the corresponding signal. Shuffling the added blocks removes the gain. The gain persists after selected label-associated proteins are removed. These controls link the improvement to cell-matched biological content. We also build and release CellSpectrum, a broader benchmark spanning eight sources and seven tasks with paired modalities. The gains hold there across tasks, tissues, and modalities. These results show that a better cell representation can improve single-cell reasoning without retraining the model. Code and CellSpectrum are publicly available at https://github.com/LiZaiyuan0619/Cell-Lens.

## Introduction

Large language models are becoming general tools across science (Brown et al. 2020; Guo et al. 2025). They read experimental data, follow written instructions, and reason over scientific questions alongside specialized tools (Wei et al. 2022; Kojima et al. 2022). Single-cell biology is now one of these areas, because its measurements are large and detailed (Theodoris et al. 2023; Cui et al. 2024). For example, researchers already use an LLM to read a cell and name its type (Hou and Ji 2024).

The standard way to show a cell to an LLM is the cell-sentence, a short list of its top genes ranked by expression (Levine et al. 2024). Most current single-cell work adopts it (Rizvi et al. 2025; Chen and Zou 2024). But this list gives only a partial picture, and it can miss the evidence that defines a cell’s identity. Ranking pushes a defining marker below the cutoff, so it never enters the list (Xiao et al. 2025). Signals from other measurements are left out from the start. The model then reasons from what remains.

When the input is this limited, the representation shapes how well the model can reason (Liu et al. 2024; Sclar et al. 2024). A cell may be measured along several axes, yet its LLM interface often exposes only ranked RNA. Here, we introduce Cell-Lens, a training-free representation that preserves available paired measurements and organizes them as typed blocks (Figure 1). It also adds marker-based summaries of cross-modal links, assay-observed zeros, and coordinated programs. Together, these blocks expose evidence that the cell-sentence discards. For example, a defining marker such as CD8A often ranks out of the list, and Cell-Lens makes it visible to the model again. The design is general. It extends to other modalities, other single-cell tasks, and many cells read at once.

**Figure 1:**
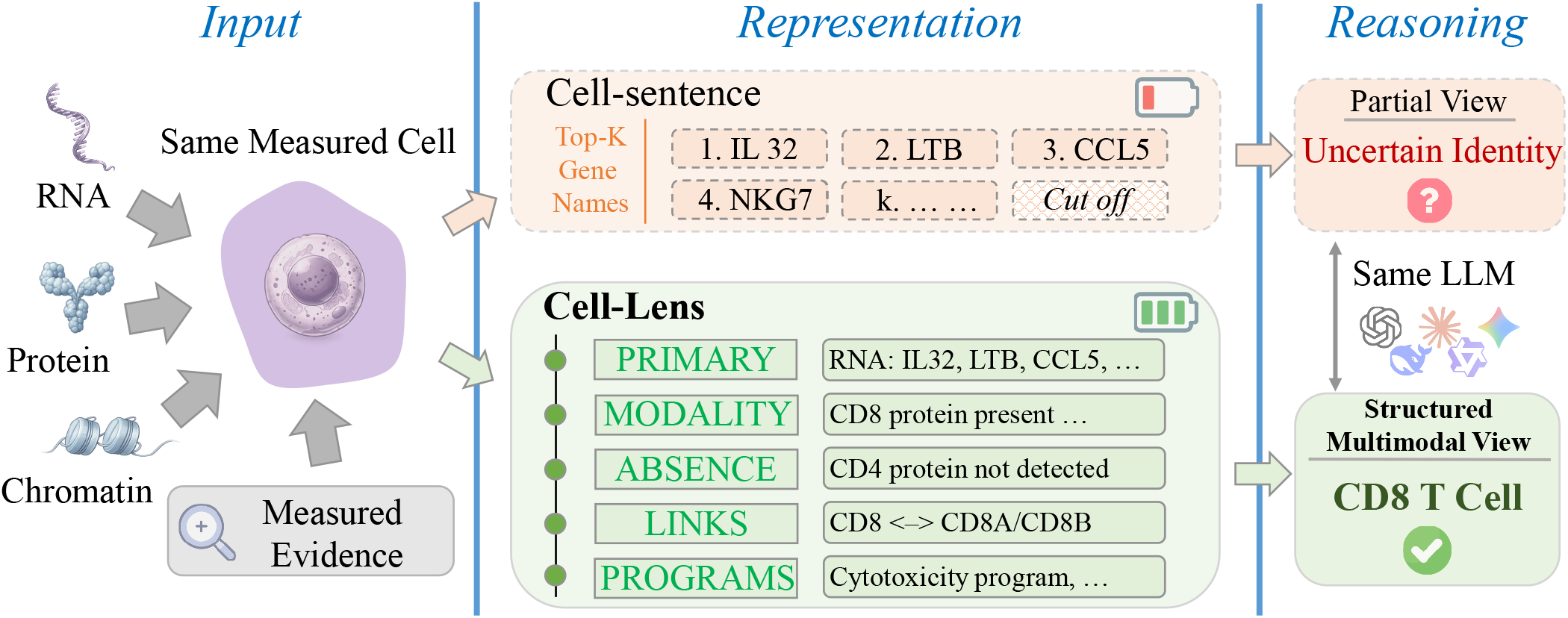
Overview of Cell-Lens. A conventional cell-sentence exposes only the top-ranked RNA genes and may omit cell-defining evidence. Cell-Lens presents paired measurements and marker-based summaries as typed blocks.

We evaluate Cell-Lens on four public benchmarks, CellVerse (Zhang et al. 2025), SOAR (Liu et al. 2025), CellPuzzles (Fang et al. 2026), and SC-Arena (Zhao et al. 2026), across five leading model families, GPT, Claude, Gemini, Qwen (Yang et al. 2025), and DeepSeek (Xu et al. 2026a). Cell-Lens raises accuracy on cell typing and on broader single-cell tasks. Ablations trace the gain to the signal we add, not to the longer input. But the public benchmarks are limited. Their released LLM inputs expose only one modality, and they cover few tasks and tissues. We therefore build and release CellSpectrum, a broader single-cell benchmark. It draws from eight sources across healthy and diseased tissue, pairs RNA with protein and with chromatin, and adds tasks the public sets do not reach. The gain holds on every task there, and it also holds on multi-cell tasks. Weaker models gain the most, so the spread between models narrows. At the same time, reported token usage falls. Representation, then, can matter as much as model choice.

We make four contributions.

1. We show that serializing a cell into a cell-sentence loses information by construction, and that this loss limits LLM reasoning about single cells.
2. We introduce Cell-Lens, a training-free typed-block representation of paired measurements and marker-based summaries.
3. We build and release CellSpectrum, a benchmark spanning eight sources, seven tasks, and paired modalities. It extends evaluation beyond the limited modalities and task types covered by existing benchmarks.
4. We find that Cell-Lens raises accuracy across benchmarks, tasks, and model families, while reducing API-reported token usage across four measured models. Controls show that the gain requires cell-matched content. It persists after selected label-associated proteins are removed.

### Related Work

Large language models reason well from given examples (Brown et al. 2020; Wei et al. 2022; Kojima et al. 2022; Guo et al. 2025). Their accuracy is not set by the model alone. The input shapes it too, for example whether the data is presented in a structured form (Liu et al. 2024). A different prompt format or example order changes the result (Lu et al. 2022; Sclar et al. 2024; Min et al. 2022). The information is the same, but the answer is not. This sensitivity is especially relevant to structured scientific objects, because their text encoding is the model’s only input. For molecules, the string encoding changes how well the model reasons (Yan et al. 2026; Edwards et al. 2022). For graphs, the text encoding decides which problems the model can solve (Xu et al. 2026b; Fatemi, Halcrow, and Perozzi 2024; Wang et al. 2023). A single cell is exactly such an object. We study how to put richer cellular evidence into text for an LLM.

Single-cell work turns a cell into a cell-sentence, a list of its top genes ranked by expression (Levine et al. 2024). This interface has been scaled up and reused as a general input format (Rizvi et al. 2025; Chen and Zou 2024). Even the choice of genes for the list changes the accuracy (Xiao et al. 2025). Recent benchmarks adopt the cell-sentence to test cell typing and reasoning (Zhang et al. 2025; Fang et al. 2026). Other benchmarks use ontology-distance scoring and knowledge-augmented evaluation (Liu et al. 2025; Zhao et al. 2026). All of them keep the ranked gene list as the interface, and that interface is coarse. It drops the other modalities, markers with assay-observed zero values, and the magnitude. Multimodal single-cell analysis has already established the value of paired assays (Hao et al. 2021). Cell-Lens addresses a different bottleneck by making this evidence readable to frozen LLMs. It retains the ranked list and adds paired measurements and marker-based summaries. Typed blocks keep these evidence types explicit.

Most single-cell work with LLMs retrains the model on omics data (Theodoris et al. 2023; Cui et al. 2024). This foundation-model line has grown to larger corpora and new architectures (Yang et al. 2022; Hao et al. 2024). Some train a dedicated cell language model or reason across many samples at once (Wen et al. 2024; Sypetkowski et al. 2026). A second line keeps the model unchanged and annotates at the cluster level by reading a marker list (Hou and Ji 2024). Classical annotators map each cell to a reference or a trained classifier, without a language interface (Aran et al. 2019; Xu et al. 2021). Cell-Lens addresses the complementary question of how to present a measured cell to a frozen LLM. It keeps the model unchanged and redesigns its per-cell text interface. This setting isolates a representation bottleneck that conventional annotation pipelines do not test.

## Method

### The Cell-Sentence

A single cell is a high-dimensional measurement along several axes. Modern assays often record two modalities at once, RNA and protein in CITE-seq (Stoeckius et al. 2017), or chromatin and protein in ASAP-seq (Mimitou et al. 2021). To query a frozen LLM, a benchmark serializes the cell into a cell-sentence (Levine et al. 2024), the top-*K* genes by expression in rank order,

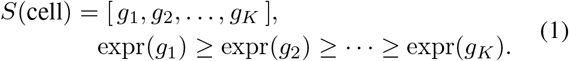

The cell-sentence therefore states the identity and rank of the most expressed genes. The benchmark sets *K*, and everything ranked below it is dropped. Whatever the assay measured outside expression is dropped with it. The ranked identity is what the model reads.

A ranked gene list thus leaves out three things, the other modalities the assay measured, markers with assay-observed zero values, and the expression magnitude. An LLM cannot recover the first from the cell-sentence, because a protein readout is not a function of the gene names. It cannot recover the second either, since a gene absent from the list looks the same as one with an assay-observed zero. The third is available in principle, but it sits close to Poisson noise (Sarkar and Stephens 2021; Tang et al. 2023).

### The Cell-Lens Representation

We start from what defines a cell’s identity, the signals measured along its several axes. Our question is how a language representation should present them. A faithful representation makes this identity visible in the text, so the model can read it directly. It should stay ordinary typed text, generated from the measurement and read by the model without any conversion. Ranking alone hides part of that identity. A marker can define a cell type through its protein, while its gene sits below the top-*K* cutoff and never enters the cell-sentence (Figure 4).

Concretely, Cell-Lens represents each cell as a single typed block (Figure 1):

~~~
[CELL assay=CITE-seq]
   PRIMARY: <expression axis: gene names,
      expression-ranked>
   MODALITY: <other measured modalities:
      surface proteins, chromatin, …;
      one entry each, identity-ranked>
   ABSENCE: <prespecified panel markers
      not detected on the measured axis>
   LINKS: <cross-modal identity:
      CD8(protein)-CD8A/CD8B(gene);
      CD3(protein)-CD3D/E/G(gene); …>
   PROGRAMS: <lineage or state programs,
      with supporting-marker counts>
[/CELL]
~~~

The header names the assay, and the assay sets which axes the cell carries. The block then reads from top to bottom, one axis at a time. PRIMARY gives the expression axis, its gene names in rank order. This axis carries the identity the model already reads well, and the blocks below give the axes it cannot. MODALITY states the other measured modalities, the surface proteins in CITE-seq or the chromatin in ASAP-seq. It takes one entry per modality, and holds as many as the assay measures. No ordering of the gene names can produce these entries. MODALITY therefore brings evidence the text never carried. Within each modality the entries are ordered by identity. The magnitude is left out, since it is as noisy as the RNA counts (Lopez et al. 2018).

ABSENCE lists prespecified panel markers with assay-observed zero values on the same axis. These zeros do not establish biological absence. They do, however, differ from nonzero markers omitted by the top-*K* cutoff. ABSENCE preserves this distinction. LINKS states the cross-modal correspondence for a shared identity, so CD8 protein and the CD8A gene stand for one signal. The model then reads the two modalities as a single identity, not as two unrelated names. PROGRAMS summarizes lineage or state evidence with prespecified marker sets. It names each supported program and reports its detected-marker count. This summary makes distributed evidence readable.

PRIMARY contains the plain cell-sentence. The standard baseline therefore sits inside this representation rather than beside it. The blocks are plain text generated from the measurement, so the model itself runs unchanged.

### Construction

Cell-Lens starts from the cell’s paired measurements. The cell barcode joins the modalities, so every block describes the same cell. Fixed rules then select entries and construct the marker-based summaries. The process requires no training or model call.

PRIMARY and MODALITY come from the cell’s own numbers alone. Each orders its entries by the quantity its axis reports. On the expression axis that quantity is the transcript count. On surface protein it is normalized abundance, and on chromatin it is TF-IDF weighted gene activity. A fixed cut on every axis keeps the block short enough to read. The length is set in advance and never tuned per cell. On the expression axis here it is the same *K* the baseline uses, so both conditions read the same genes.

The remaining three blocks use biological references frozen before evaluation. ABSENCE checks a prespecified lineage panel for assay-observed zeros. PROGRAMS aggregates detected markers with prespecified sets. LINKS uses fixed protein-gene correspondences. Block construction does not access the target annotation or model output.

### Generality

The representation is not tied to any one assay or task. The block types instantiate on whatever axes an assay measures, so the same blocks carry other modalities and serve other single-cell tasks. A new axis or program enters as one more typed entry, and the block types stay the same.

The representation can encode identity signals from any measured axis as typed entries. The representation therefore applies wherever a single cell is measured beyond a ranked gene list. The tasks we report are the ones current benchmarks support, and the same blocks are ready for the assays those benchmarks do not yet cover.

## Experiments

### Experimental Setup

We evaluate frozen frontier LLMs through their provider APIs with no weight change and no local training. We hold the cell and model fixed in every comparison, while changing the input interface. This paired design isolates how the text interface affects a frozen LLM. We cover the major model families, including GPT, Claude, Gemini, Qwen (Yang et al. 2025), and DeepSeek (Xu et al. 2026a), and use at least two recent frontier models in each. The exact models are named in the tables. Throughout, Baseline uses the released cell-sentence, whereas Ours adds available paired measurements and marker-based summaries in typed blocks. We report accuracy in points and the gain of Ours over Baseline as Δ. Each model is queried zero-shot, with one system message and one user message and no worked examples. Decoding is greedy at temperature 0, except on CellPuzzles, which follows its official temperature-1 setup. For each task, we select one fixed class-balanced subset and use the same cells in every paired comparison. Provider-reported token counts are paired within each model and API, and total usage combines input, output, and reasoning tokens.

We build the evaluation on four public single-cell benchmarks, CellVerse (Zhang et al. 2025), SOAR (Liu et al. 2025), CellPuzzles (Fang et al. 2026), and SC-Arena (Zhao et al. 2026). Each benchmark serializes cells as cell-sentences and retains its official metric. CellVerse uses exact match, SOAR and SC-Arena use ontology-distance scores, and CellPuzzles uses batch accuracy. The CellVerse cell-typing result draws on three independent CITE-seq sources, Mimitou (GSE156478) (Mimitou et al. 2021), a 10x PBMC set with TotalSeq-B, and Hao 2021 (GSE164378) (Hao et al. 2021). Together these benchmarks span a range of tissues and task types, and they cover much of what current work asks of an LLM on single-cell data.

### The CellSpectrum Benchmark

Yet these four benchmarks share three limitations. Each provides a single-modality text interface, covers a narrow set of tasks, and drops cross-modal barcodes during serialization. We therefore build and release CellSpectrum, which retains paired measurements and cross-modal barcodes.

CellSpectrum draws from eight sources spanning blood, bone marrow, and solid organs. These include healthy bone marrow (BMMC, GSE194122) (Lance et al. 2022), diseased lymphoid tissue (T-ALL, GSE248287) (Wiggers et al. 2025), heart tissue (GSE217494) (Amrute et al. 2024), and genotype-labeled real doublets (GSE96583) (Kang et al. 2018). CellSpectrum retains the labels defined by each source or task protocol, while Cell-Lens blocks are constructed without accessing them. The bone marrow and lymphoid samples test transfer across tissue and disease. The heart is a solid organ, the furthest from blood here, and it tests whether the method breaks there. The doublets carry a genotype label, so they test the calls against independent ground truth. CellSpectrum pairs RNA with protein and with chromatin, and it keeps the cross-modal barcode the public sets discard. On top of cell typing, it adds task types the public benchmarks do not reach, including doublet quality control (Wolock, Lopez, and Klein 2019; McGinnis, Murrow, and Gartner 2019), cell-cycle detection (Tirosh et al. 2016), and de-novo typing. Together CellSpectrum spans a broader range of tissues, modalities, and task types, and it allows a more complete evaluation of what an LLM can do on single-cell data.

### Main Results

#### Cell-Lens lifts accuracy across all four public benchmarks and six models (Table 1)

The gain is stable in most settings. The cell and frozen model remain fixed, while Cell-Lens changes the available evidence and its organization. The gains therefore demonstrate the value of the Cell-Lens representation as a whole.

**Table 1:** Main comparison across four published benchmarks, one per row. B is Baseline, O is Ours, and Δ is the gain of O over B in points. CellVerse is split into its three CITE sources (10x-PBMC, GSE156478, GSE164378) and the ASAP chromatin subtask. Each benchmark keeps its own official metric, so columns are not on one scale.

|  |  | GPT-5.4-mini |  |  | Qwen3.7-Plus |  |  | DeepSeek-V4-Flash |  |  | Gemini-3.5-Flash |  |  | GPT-5.6-Sol |  |  | Claude-Opus-4-8 |  |  |
| --- | --- | --- | --- | --- | --- | --- | --- | --- | --- | --- | --- | --- | --- | --- | --- | --- | --- | --- | --- |
| Benchmark | | B | O | $\Delta$ | B | O | $\Delta$ | B | O | $\Delta$ | B | O | $\Delta$ | B | O | $\Delta$ | B | O | $\Delta$ |
| CellVerse | 10x-PBMC | 67.4 | 81.7 | +14.3 | 67.4 | 92.0 | +24.6 | 65.4 | 91.7 | +26.3 | 64.9 | 92.0 | +27.1 | 72.3 | 89.1 | +16.8 | 72.9 | 91.7 | +18.8 |
|  | GSE156478 | 59.4 | 68.0 | +8.6 | 52.3 | 74.0 | +21.7 | 55.7 | 72.0 | +16.3 | 52.6 | 75.1 | +22.5 | 59.4 | 76.0 | +16.6 | 60.9 | 78.0 | +17.1 |
|  | GSE164378 | 61.7 | 72.3 | +10.6 | 60.3 | 72.3 | +12.0 | 61.1 | 76.3 | +15.2 | 56.6 | 74.3 | +17.7 | 66.3 | 78.0 | +11.7 | 66.9 | 77.7 | +10.8 |
|  | ASAP | 44.3 | 63.8 | +19.5 | 39.0 | 59.0 | +20.0 | 48.1 | 63.3 | +15.2 | 37.7 | 60.9 | +23.2 | 45.4 | 72.3 | +26.9 | 46.3 | 64.6 | +18.3 |
| SOAR |  | 42.2 | 54.8 | +12.6 | 39.6 | 47.0 | +7.4 | 33.8 | 38.6 | +4.8 | 46.4 | 57.8 | +11.4 | 41.2 | 53.0 | +11.8 | 42.8 | 55.0 | +12.2 |
| CellPuzzles |  | 28.3 | 30.8 | +2.5 | 13.3 | 30.8 | +17.5 | 21.7 | 31.7 | +10.0 | 34.2 | 45.0 | +10.8 | 30.8 | 38.3 | +7.5 | 35.0 | 36.7 | +1.7 |
| SC-Arena |  | 30.9 | 33.2 | +2.3 | 32.3 | 34.4 | +2.1 | 27.8 | 31.7 | +3.9 | 41.1 | 43.7 | +2.6 | 28.7 | 30.9 | +2.2 | 24.8 | 32.5 | +7.7 |

#### The gain extends to more models and narrows the gap between them (Table 2)

Beyond the six models above, the same representation helps four further models. The effect is not tied to a particular model. Cell-Lens also brings the models closer together. Under the raw cell-sentence they span a wide range, and after the change they converge near the top. The weakest baseline model gains the most, reducing the performance spread across models.

**Table 2:** Convergence across four models not already in Table 1, on CITE (mean of the three sources), ASAP, and CellPuzzles.

| Benchmark |  | Qwen3.6<br>Flash | DeepSeek<br>V4-Pro | GPT<br>5.5 | Claude<br>Opus-4-7 |
| --- | --- | --- | --- | --- | --- |
| CITE | Base | 51.2 | 63.0 | 67.0 | 59.7 |
|  | Ours | 78.3 | 83.5 | 82.2 | 75.9 |
| | $\Delta$ | +27.1 | +20.5 | +15.2 | +16.2 |
| ASAP | Base | 36.2 | 54.3 | 40.5 | 41.4 |
|  | Ours | 51.0 | 66.2 | 55.7 | 57.1 |
| | $\Delta$ | +14.8 | +11.9 | +15.2 | +15.7 |
| CellPuzzles | Base | 11.7 | 30.0 | 37.5 | 31.2 |
|  | Ours | 19.2 | 36.8 | 40.8 | 34.8 |
| | $\Delta$ | +7.5 | +6.8 | +3.3 | +3.6 |

#### Cell-Lens improves accuracy while reducing API-reported token usage (Figure 2)

Across every model shown, reasoning-token usage falls by 22% to 31%, while total token usage falls by 4% to 10%. Accuracy rises at the same time. The gain also persists across all tested reasoning-effort settings and three model families (Figure 3). Together, these results show that Cell-Lens improves the observed accuracytoken tradeoff.

**Figure 2:**
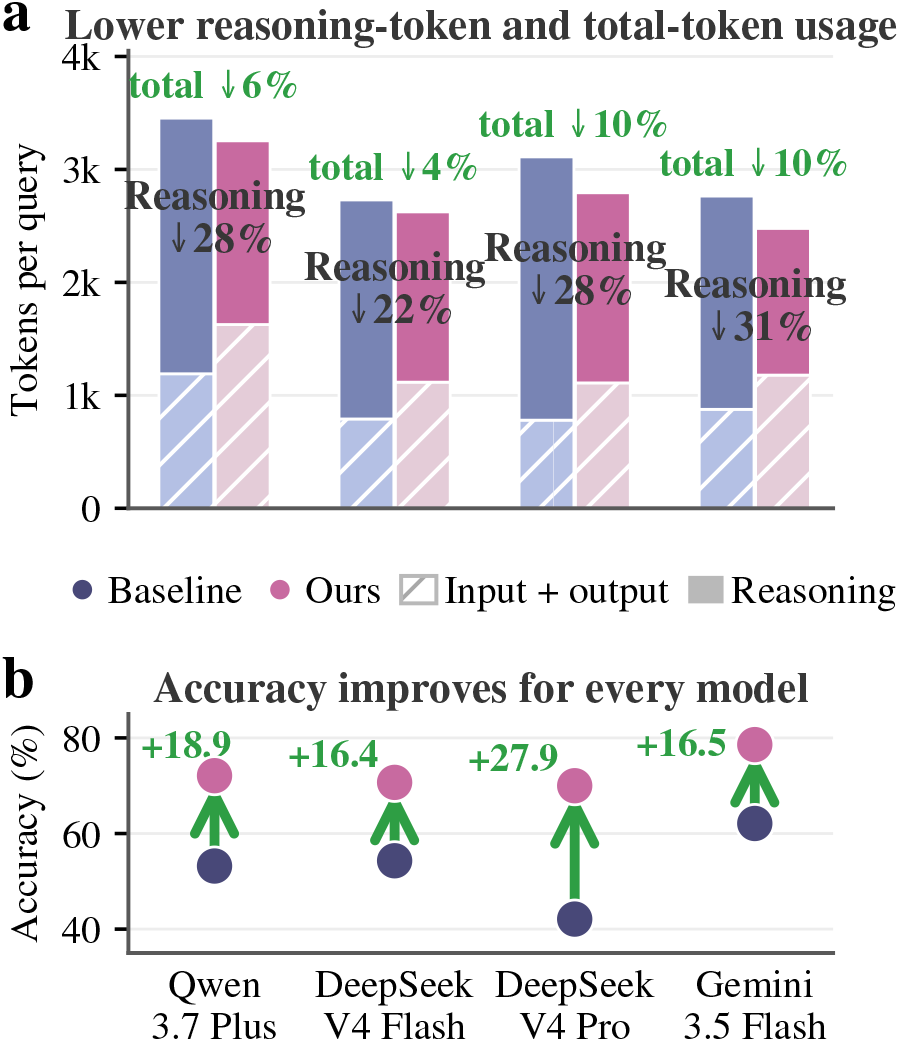
Cell-Lens reduces API-reported token usage while improving accuracy. Panel (a) reports reasoning-token and total-token usage per query. Panel (b) shows accuracy gains across all four models.

**Figure 3:**
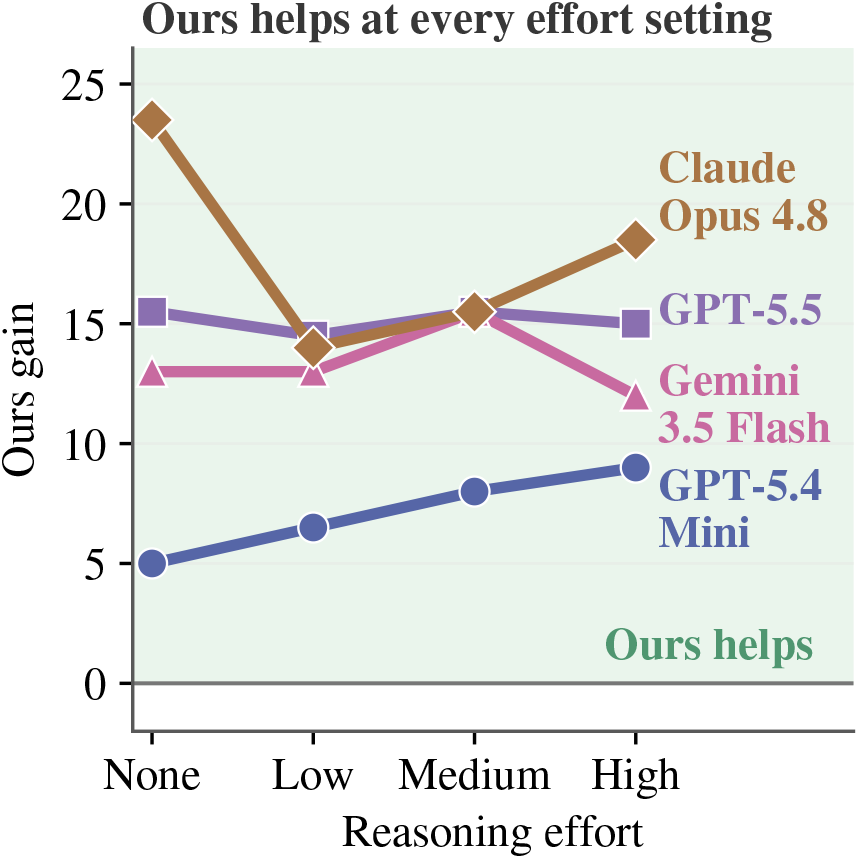
Cell-Lens improves accuracy across models at every tested reasoning-effort setting. At each fixed setting, this gain requires no increase in configured effort.

**Figure 4:**
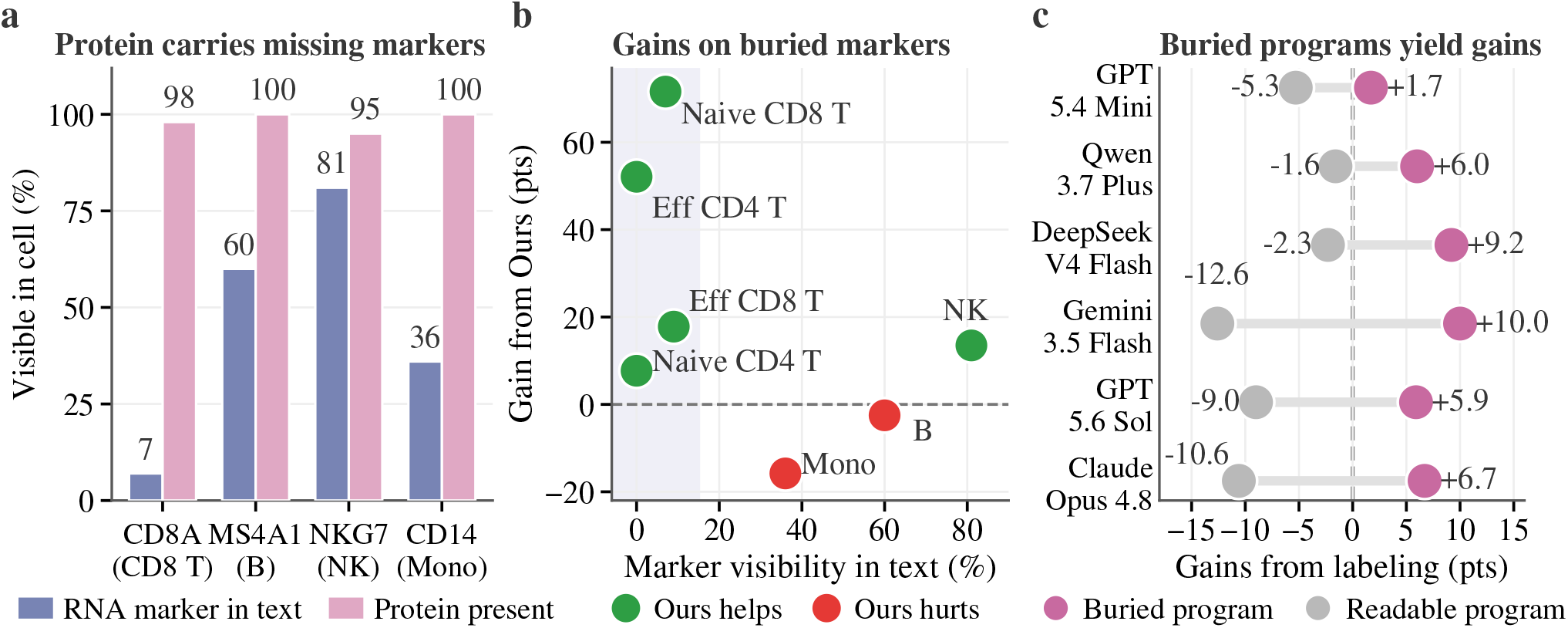
The gain has a mechanism. (a) A defining RNA marker is often ranked out of the cell-sentence while its protein is present. (b) Per-cell gains are largest for cell types whose defining markers are buried in the text. They are small or negative for most cell types whose markers are already visible. (c) A marker-based summary of a buried cell-cycle program raises accuracy across models. A summary of an already-readable high-mitochondrial-content program lowers accuracy.

### Mechanism

#### Cell-Lens exposes biologically informative signals omitted from the cell-sentence (Figure 4)

A defining marker is often ranked out of the cell-sentence while another measured modality still carries it. The clearest case is CD8A, the marker of CD8 T cells. It appears in only 7% to 9% of their cell-sentences, while the protein carries the same identity at 93% to 98%. A model that reads only the cell-sentence guesses these cells at chance. Across cell types, gains are largest when defining markers are absent from the cell-sentence.

#### Ablations link the gain to cell-matched content (Table 3)

A shuffle swaps a cell’s block for another cell’s. Accuracy then falls below the baseline. Equal-length but mismatched text is therefore insufficient. Removing selected label-associated proteins (CD3/4/8/19/14/56/45RA/45RO) still leaves a clear gain. These direct markers do not explain the full improvement. Dropping the protein values changes nothing. The ranked identities carry the signal, not their magnitudes. Together, the controls tie the gain to cell-matched biological content.

**Table 3:** Controls on the CITE gain, across three sources and six models. Shuffle takes the block from another cell, and ablate removes selected label-associated proteins. **Bold** marks the best value in each row.

| Model | Source | Baseline | Ours | Shuffle | Ablate |
| --- | --- | --- | --- | --- | --- |
| GPT-5.4-mini | 10x-PBMC | 67.4 | **81.7** | 53.1 | 72.0 |
|  | GSE156478 | 59.4 | **68.0** | 37.4 | 61.4 |
|  | GSE164378 | 61.7 | **72.3** | 54.3 | 63.7 |
| Qwen3.7-Plus | 10x-PBMC | 67.4 | **92.0** | 20.6 | 73.7 |
|  | GSE156478 | 52.3 | **74.0** | 22.6 | 62.3 |
|  | GSE164378 | 60.3 | **72.3** | 39.4 | 59.7 |
| DeepSeek-V4-Flash | 10x-PBMC | 65.4 | **91.7** | 18.8 | 75.1 |
|  | GSE156478 | 55.7 | **72.0** | 17.7 | 65.7 |
|  | GSE164378 | 61.1 | **76.3** | 33.1 | 57.7 |
| Gemini-3.5-Flash | 10x-PBMC | 64.9 | **92.0** | 31.1 | 74.6 |
|  | GSE156478 | 52.6 | **75.1** | 30.3 | 62.3 |
|  | GSE164378 | 56.6 | **74.3** | 45.7 | 59.1 |
| GPT-5.6-Sol | 10x-PBMC | 72.3 | **89.1** | 54.3 | 73.4 |
|  | GSE156478 | 59.4 | **76.0** | 37.1 | 65.4 |
|  | GSE164378 | 66.3 | **78.0** | 55.1 | 65.4 |
| Claude-Opus-4-8 | 10x-PBMC | 72.9 | **91.7** | 35.1 | 76.0 |
|  | GSE156478 | 60.9 | **78.0** | 29.7 | 68.6 |
|  | GSE164378 | 66.9 | **77.7** | 52.0 | 64.0 |

#### A de-novo comparison separates matched protein evidence from marker-name annotation (Table 4)

The public benchmarks drop the barcode that links a cell across modalities. Their official sets therefore cannot support a modality-versus-label contrast. We run that contrast on a de-novo typing task we build ourselves. Ours adds matched protein evidence, whereas Label names markers already present in the RNA input. Across all six models, matched protein evidence improves accuracy, while Label ties or reduces it.

**Table 4:** De-novo typing. B uses the RNA input, O adds matched protein evidence, and L names markers already visible in B. Each Δ is against B in points.

| Model | B | O | $\Delta_O$ | L | $\Delta_L$ |
| --- | --- | --- | --- | --- | --- |
| DeepSeek-V4-Flash | 32.2 | 79.7 | +47.5 | 32.2 | +0.0 |
| Qwen3.7-Plus | 49.0 | 93.0 | +44.0 | 49.0 | +0.0 |
| Gemini-3.5-Flash | 62.6 | 69.6 | +7.0 | 43.1 | -19.5 |
| GPT-5.4-mini | 65.8 | 76.0 | +10.2 | 43.1 | -22.7 |
| Claude-Opus-4-8 | 70.6 | 74.6 | +4.0 | 68.5 | -2.1 |
| GPT-5.6-Sol | 76.2 | 78.3 | +2.1 | 53.7 | -22.5 |

### Generalization

#### The gain of Cell-Lens extends to task types the public benchmarks do not cover (Table 6)

The gain so far held across ten models, and we now test whether it also crosses tasks, tissues, and modalities. Cell-Lens helps on doublet detection, cell-cycle detection, and de-novo typing. It also holds across tissue and disease, from healthy bone marrow (BMMC) to a diseased lymphoid sample (T-ALL).

#### On real genotype-labeled doublets, Cell-Lens improves detection accuracy and reports more complete lineage assignments (Figure 5)

The fraction with both reference lineages named rises from Baseline to Ours, making the output easier to verify.

**Figure 5:**
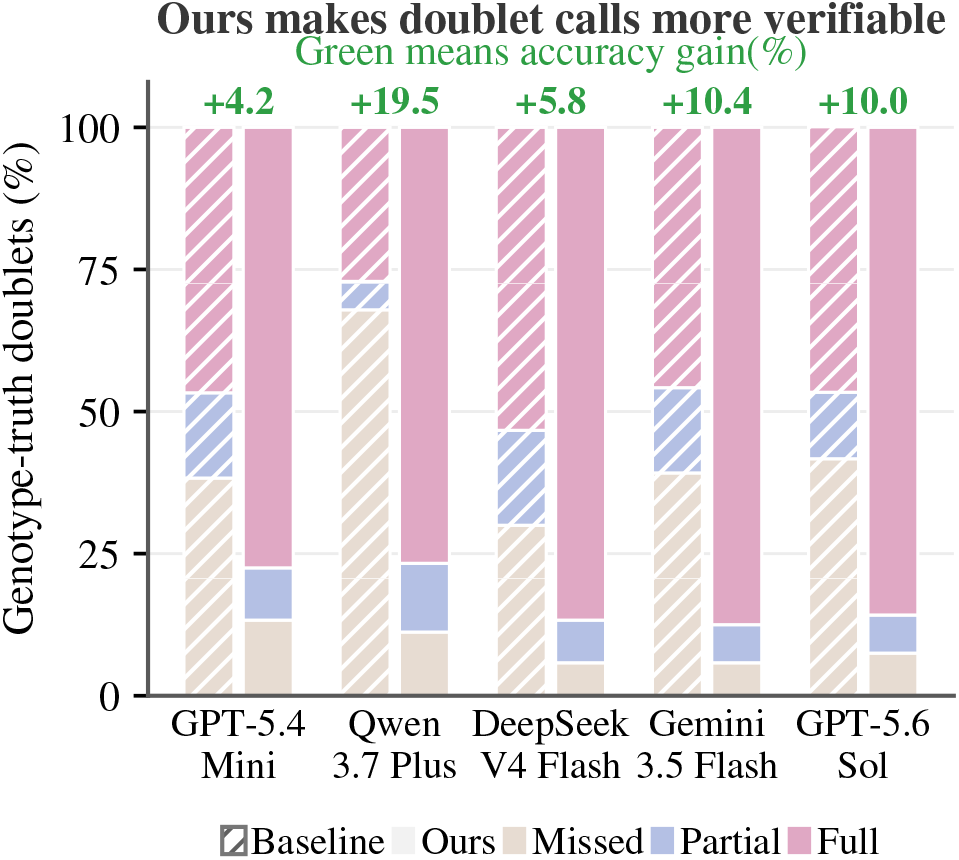
Real genotype-labeled doublets. Stacked bars show whether a true doublet is missed or whether one or both reference lineages are named. Cell-Lens increases both the “both named” fraction and detection accuracy (line, right axis).

#### Cell-Lens extends to additional modalities, such as chromatin (Table 5)

The results so far paired RNA with protein. SOAR pairs RNA with chromatin and shows gains across models. A CellSpectrum multiome task shows a similar pattern. Positive gains on both benchmarks extend the results beyond protein-paired data.

**Table 5:**
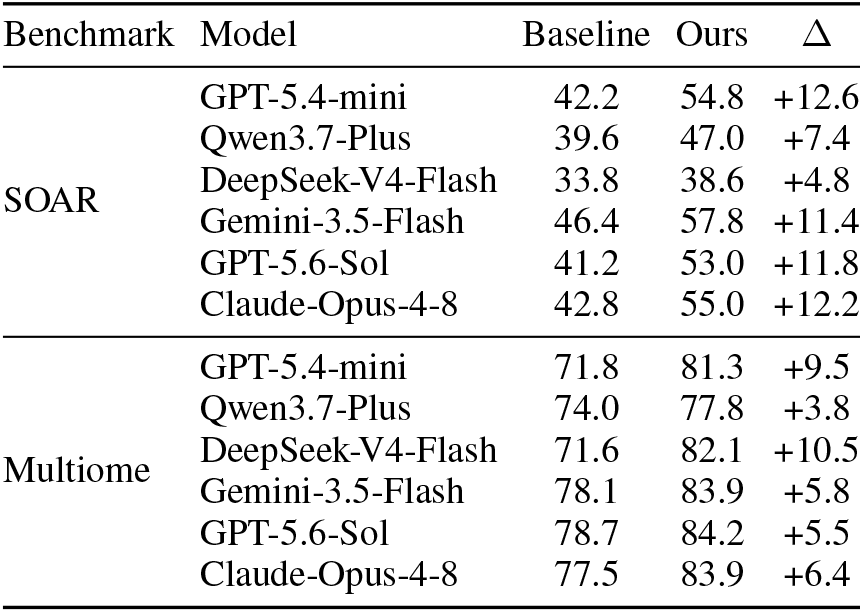
Chromatin as the modality paired with RNA. Two benchmarks pair RNA with chromatin, the public SOAR and a self-built multiome task.

| Benchmark | Model | Baseline | Ours | $\Delta$ |
| --- | --- | --- | --- | --- |
| SOAR | GPT-5.4-mini | 42.2 | 54.8 | +12.6 |
|  | Qwen3.7-Plus | 39.6 | 47.0 | +7.4 |
|  | DeepSeek-V4-Flash | 33.8 | 38.6 | +4.8 |
|  | Gemini-3.5-Flash | 46.4 | 57.8 | +11.4 |
|  | GPT-5.6-Sol | 41.2 | 53.0 | +11.8 |
|  | Claude-Opus-4-8 | 42.8 | 55.0 | +12.2 |
| Multiome | GPT-5.4-mini | 71.8 | 81.3 | +9.5 |
|  | Qwen3.7-Plus | 74.0 | 77.8 | +3.8 |
|  | DeepSeek-V4-Flash | 71.6 | 82.1 | +10.5 |
|  | Gemini-3.5-Flash | 78.1 | 83.9 | +5.8 |
|  | GPT-5.6-Sol | 78.7 | 84.2 | +5.5 |
|  | Claude-Opus-4-8 | 77.5 | 83.9 | +6.4 |

**Table 6:**
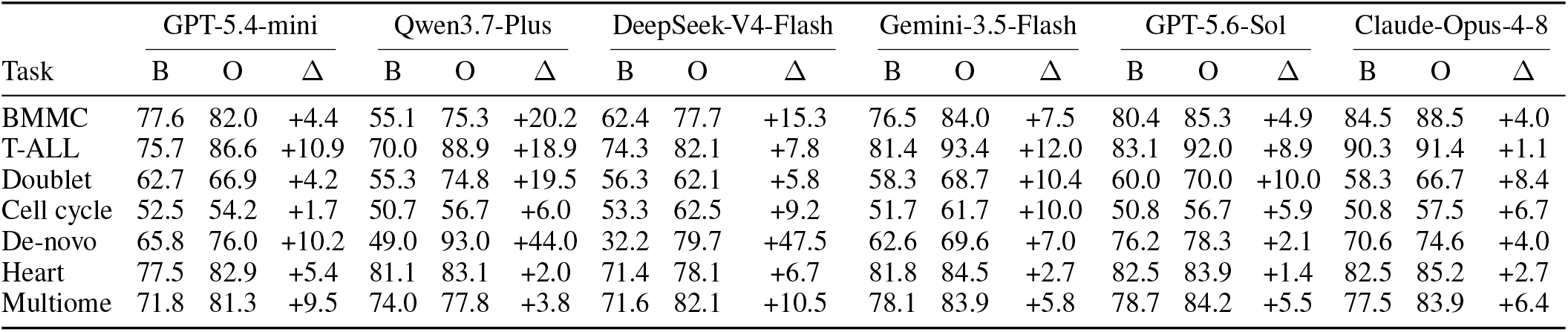
The CellSpectrum benchmark, seven single-cell tasks across six models. B is Baseline, O is Ours, and Δ is the gain of O over B in points.

#### Per-cell blocks improve multi-cell accuracy across all six models (Table 7)

This comparison holds cellular content fixed and changes only its organization. Flat text can obscure cell boundaries, whereas per-cell blocks preserve them. The gains range from 4.7 to 12.9 points.

**Table 7:** The multi-cell task in CellSpectrum, where one prompt holds many cells. Both conditions contain the same cells and evidence. Flat concatenates them as one text, whereas Ours preserves one typed block per cell. Δ is the gain of Ours over Flat in points.

| Model | Flat | Ours | $\Delta$ |
| --- | --- | --- | --- |
| GPT-5.4-mini | 56.9 | 62.9 | +6.0 |
| Qwen3.7-Plus | 64.3 | 69.0 | +4.7 |
| DeepSeek-V4-Flash | 55.2 | 68.1 | +12.9 |
| Gemini-3.5-Flash | 56.7 | 67.3 | +10.6 |
| GPT-5.6-Sol | 58.1 | 67.6 | +9.5 |
| Claude-Opus-4-8 | 62.4 | 70.2 | +7.8 |

## Conclusion

On single-cell typing, input representation often constrains model performance. We introduce Cell-Lens, a training-free representation of paired measurements and marker-based summaries. Typed blocks keep modalities, cross-modal links, and coordinated programs explicit. This raises accuracy in cell typing and broader single-cell tasks. It also narrows performance differences between models while reducing API-reported token usage. The gains hold across tasks and tissues, and on both protein and chromatin. Under matched content, per-cell blocks also outperform flat text across all six models. Control experiments link these gains to biologically matched signals. We also release CellSpectrum, a benchmark for a more complete test of single-cell reasoning. Cell-Lens complements dedicated annotation and multimodal analysis pipelines by making richer measured evidence accessible to frozen LLMs. Future work could extend the representation to additional modalities and automate its construction. End-to-end evaluation in real analysis workflows is another important next step. For an LLM that reads a scientific object, how the object is presented can matter as much as the choice of model.

## Acknowledgments

We thank Zhehong Ai of Shanghai Artificial Intelligence Laboratory for helpful discussions.

